# Optimized extraction method enables quantitative analysis of surface metabolite recovery for exposomics and behavioral studies

**DOI:** 10.1101/2021.08.25.457715

**Authors:** Mitchelle Katemauswa, Ekram Hossain, Zongyuan Liu, Mahbobeh Lesani, Adwaita R. Parab, Danya A. Dean, Laura-Isobel McCall

## Abstract

Workplace chemical exposures are a major source of occupational injury. Although over half of these are skin exposures, exposomics research often focuses on chemical levels in the air or in worker biofluids such as blood and urine. Until now, one limitation has been the lack of methods to quantitatively measure surface chemical transfer. Outside the realm of harmful chemicals, the small molecules we leave behind on surfaces can also reveal important aspects of human behavior. In this study, we developed a swab-based quantitative approach to determine small molecule concentrations across common surfaces. We demonstrate its utility using one drug, cyclobenzaprine, and two human-derived metabolites, carnitine and phenylacetylglutamine, on four common surfaces: linoleum flooring, plastified laboratory workbench, metal and Plexiglass. This approach enabled linear small molecule recovery and quantification of molecule abundance on workplace built environment surfaces. Overall, this method paves the way for future quantitative exposomics studies.

---

Chemical exposures are a leading cause of worker injury and occupational disease ^1^. At least 13 million workers are at risk of workplace skin chemical exposure ^1a^. Such exposures lead to cutaneous symptoms (e.g. contact dermatitis, topical allergies), or systemic manifestations, up to and including cancer ^1a^. Of the 50,000 occupational chemical exposures in 2012, 34,400 were skin exposures, though this is likely an underestimate due to under-reporting ^1^. Corrected estimates range from 250,000 to 1.25 million occupational contact skin disease cases ^1b^. Workplace chemical skin exposures cost the US economy over $1 billion per year (including treatment costs, days of work lost, reduced productivity) ^1a^ and also cause significant social and psychological burden ^1b^.

However, most studies of chemical exposures focus on respiratory occupational exposures. Indeed, while there is an OSHA list of permissible chemical occupational exposure limits, these all refer to exposure by inhalation and no such list exists for skin exposures (OSHA, NIOSH and the American Conference of Industrial Hygienists do list a few hundred of those chemicals with skin penetrability (‘skin notation’)). Within offices, retail spaces, hospitals, etc., workers are constantly touching building surfaces and objects. We and others have shown that these surfaces share chemical profiles with the people touching them ^2^, indicating chemical exchanges between building surfaces and workers and the potential for occupational chemical exposure. Occupational health studies also usually focus on the workers directly handling these known risk agents. However, our prior work has shown traces of laboratory chemicals outside of the research space ^2a^ and shared chemical signatures for research and mixed-use buildings compared to single-use office buildings ^3^, indicating that all building occupants or visitors may be at risk of exposures, and not just those handling the hazardous chemicals. Thus, there is a need to quantitatively assess chemical recovery from surfaces, to pave the way for improved analyses of chemical skin exposure risks.

Beyond occupational exposures, surface chemical traces represent an invaluable insight into building occupant behavior, enabling researchers to directly investigate interactions between humans and the built environment, independent of study participant recall bias or mis-representation ^4^. Swab-based analyses have helped characterize the interactions of people with their houses ^2a, 5^ and workplaces ^6^. However, until now, these analyses have all been qualitative ^7^. While such a qualitative approach successfully differentiates between highly divergent settings and identifies behavioral metabolites that are unique to one setting, quantitative approaches would enable improved analysis of environments or people with smaller differences in behavior, and comparisons between participants or study sites.

To address this issue, we developed an optimized method to recover small molecules from swabs, for quantitative liquid chromatography-mass spectrometry (LC-MS). We apply this method to five metabolites previously identified in our prior studies of the built environment ^2a, 5-6^, demonstrating linear recovery from multiple common built environment surfaces. This method is straightforward, and we anticipate broad utility for exposomics and behavioral studies.

## METHODS

### Materials

Carnitine (VWR), cyclobenzaprine (Cerilliant), erythromycin (Sigma), oleanolic acid (Cayman) and phenylacetylglutamine (Cayman) were prepared in 50% methanol and stored at 4°C until usage. Cotton swabs (Puritan) were soaked overnight in 50% HPLC-grade ethanol (Sigma) in LC-MS grade water (Fisher Optima), and then the soaking solution replaced with fresh 50% ethanol until use, as previously described ^6a, 7^. Chromatography solvents, including LC-MS grade water, acetonitrile and formic acid, were purchased from Fisher (Optima grade).

### Surface chemical spotting

Four surfaces common in an office or laboratory environment were selected: linoleum flooring, plastified worktop surfaces, metal, and Plexiglass. Each surface was cleaned with 50% HPLC-grade ethanol in LC-MS grade water and divided into 3” × 3” areas. 10-point dilutions of each of the standard chemicals were prepared and 50 μL of each dilution spotted at each position. All experiments were performed in triplicate.

### Quantitative swab-based recovery

Custom Eppendorf tubes for extract recovery were created by cutting approximately 3 mm from the bottom of a standard 0.5 mL Eppendorf tube (VWR). The tubes were then soaked in 70% ethanol to remove small molecule contaminants. The spotted chemicals were collected using a presoaked cotton swab to wipe the surface extensively for about 5 seconds. The cotton swab was placed inside the cut Eppendorf tube suspended in a standard 1.7 mL Eppendorf tube (Axygen, **Figure 1**), and the stem of the swab was cut off. The combined assemblage was centrifuged at 14,800 rpm for 15 minutes at 4°C, enabling the recovery of the surface metabolite extract from the swab into the bottom 1.7 mL Eppendorf.

**Figure 1.**
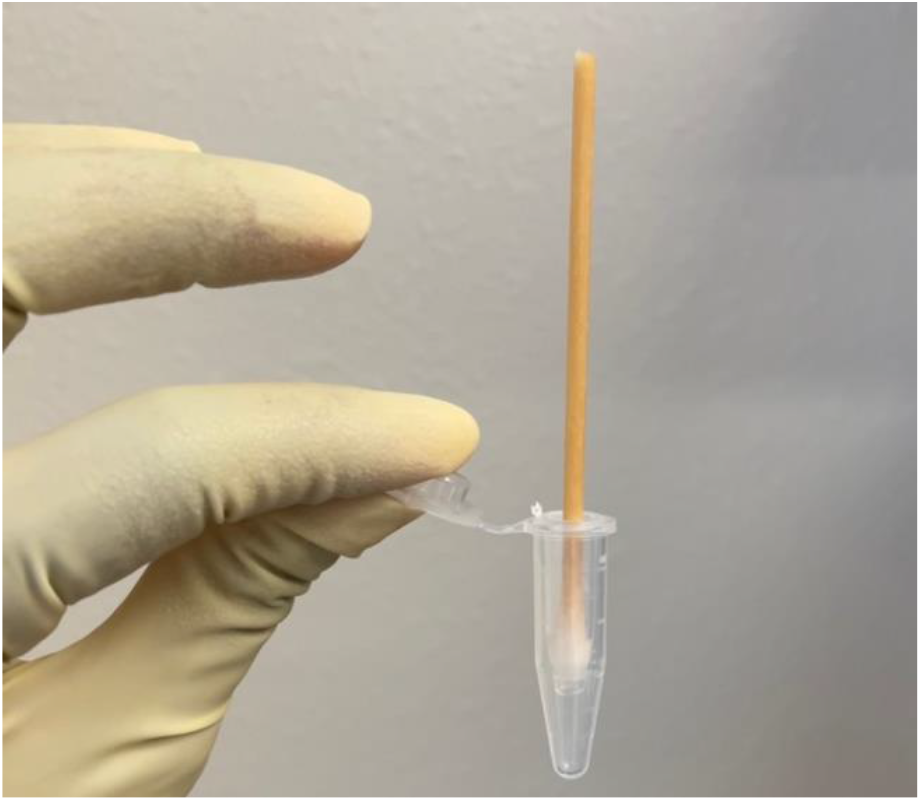
Swab and collection tube assembly.

### Implementation: workplace surface sampling

Common metal workplace samples were swabbed in the same way, including: a microwave door handle, six door handles and one stair handrail, one tap, and one water fountain button. In all cases, the surface area swabbed and the duration of swabbing matched with the standard recovery protocol.

### Sample Preparation for LC-MS

Recovered extracts were transferred to 96-well deepwell plates and dried overnight (Thermo Fisher Speedvac vacuum concentrator). Prior to LC-MS analysis, dried extracts were resuspended in 200 μL of HPLC grade 50% methanol spiked with 2 μM sulfadimethoxine as internal standard and sonicated for 15 minutes. The supernatants were then transferred to a new 96 well plate for LC-MS analysis.

### Parallel Reaction Monitoring (PRM) quantification

LC-MS data acquisition was performed using a Thermo Fisher Q Exactive Plus hybrid quadrupole-orbitrap mass spectrometer coupled to a Thermo Fisher Scientific Vanquish Flex Binary LC system. Mobile phase A was LC-MS grade water with 0.1% formic acid (Fisher Optima) and mobile phase B was LC-MS grade acetonitrile with 0.1% formic acid (Fisher Optima). LC gradient was as follows: 0-1 min, 5% B; 1-9 min, linear increase to 100% B; 9-11 min, hold at 100% B; 11-11.5 min, re-equilibrate to 5% B; 11.5-12.5 min, hold at 5% B. Flow rate was 0.5 mL/min. Samples were randomized with injections of 30 μL, with autosampler held at 10°C. A blank was run between every 12 samples. Samples were injected onto a Kinetex C18 LC column (Phenomenex; 50 × 2.1 mm, 1.7 μM particle size, 100 Å pore size) held at 40°C. All MS data acquisition was in positive mode under Parallel Reaction Monitoring (PRM). Heated electrospray ionization (HESI) source parameters were as follows: spray voltage, 3.8 kV; capillary temperature, 320 °C; sheath gas flow rate, 35; auxiliary gas flow rate, 10; sweep gas flow rate, 0; probe heater temperature, 350 °C; S-lens RF level, 50 V. PRM parameters were as follows: default charge state, 1; MS2 resolution, 17,500; AGC target, 2e5; maximum IT, 54 ms; isolation window, 1 *m/z;* stepped normalized collision energy (NCE), 20, 40, 60.

### Data processing

Raw MS files were uploaded to Skyline version 20. 1. 0. 155 ^8^. Parameters were as follows. The maximum *m/z* was 1500, and the minimum was 50. MS1 filtering was set to 70,000 resolving power at 200 *m/z* and MS/MS filtering at 17,500 resolving power at the same *m/z*. Transition list consisting of the names of precursors, fragments of interest, collision energies, and expected retention time were uploaded (see supplementary table ST-1). Peak area was exported from Skyline as comma-separated values. Plots and linear trendline analysis were performed using Microsoft Excel (Office 365).

### Data availability

Data has been deposited in MassIVE, accession numbers MSV000087739 for triplicate analysis of carnitine, cyclobenzaprine and phenylacetylglutamine; MSV000084385 for oleanolic acid and tangeritin; MSV000087746 for implementation studies.

## RESULTS AND DISCUSSION

### Linearity of recovery

Metal building surfaces had the greatest diversity of unique chemicals recovered in our prior built environment analyses, likely due to their nature as high-touch surfaces (door handles, elevator buttons, handrails, garbage cans) ^6a^. Thus, we focused our initial analyses on recovery from metal surfaces. We tested three chemicals previously detected in our built environment analyses: cyclobenzaprine, carnitine and phenylacetylglutamine ^2a, 5-6^. We anticipate higher concentrations of human- and food-derived metabolites on surfaces, so recovery of carnitine and phenylacetylglutamine was assessed in the 0.1-100 μM range. In contrast, we expect lower levels of drugs in the built environment, so recovery of cyclobenzaprine was assessed in the 0.01 nM to 0.1 μM range. Compounds were dispensed onto a metal surface at the stated concentrations and recovered using swab-based extraction 45 min after compound spotting, followed by PRM-based quantification. On metal surfaces, carnitine and phenylacetylglutamine PRM peak areas showed linearity down to our lowest tested concentration of 0.1 μM (R^2^ for linear trendline 0.9919 and 0.9997, respectively, **Figure 2AB**, purple dataset). Cyclobenzaprine showed linearity in the 0.01 nM-0.1 μM range (R^2^=0.9615, **Figure 2C**, purple dataset).

**Figure 2.**
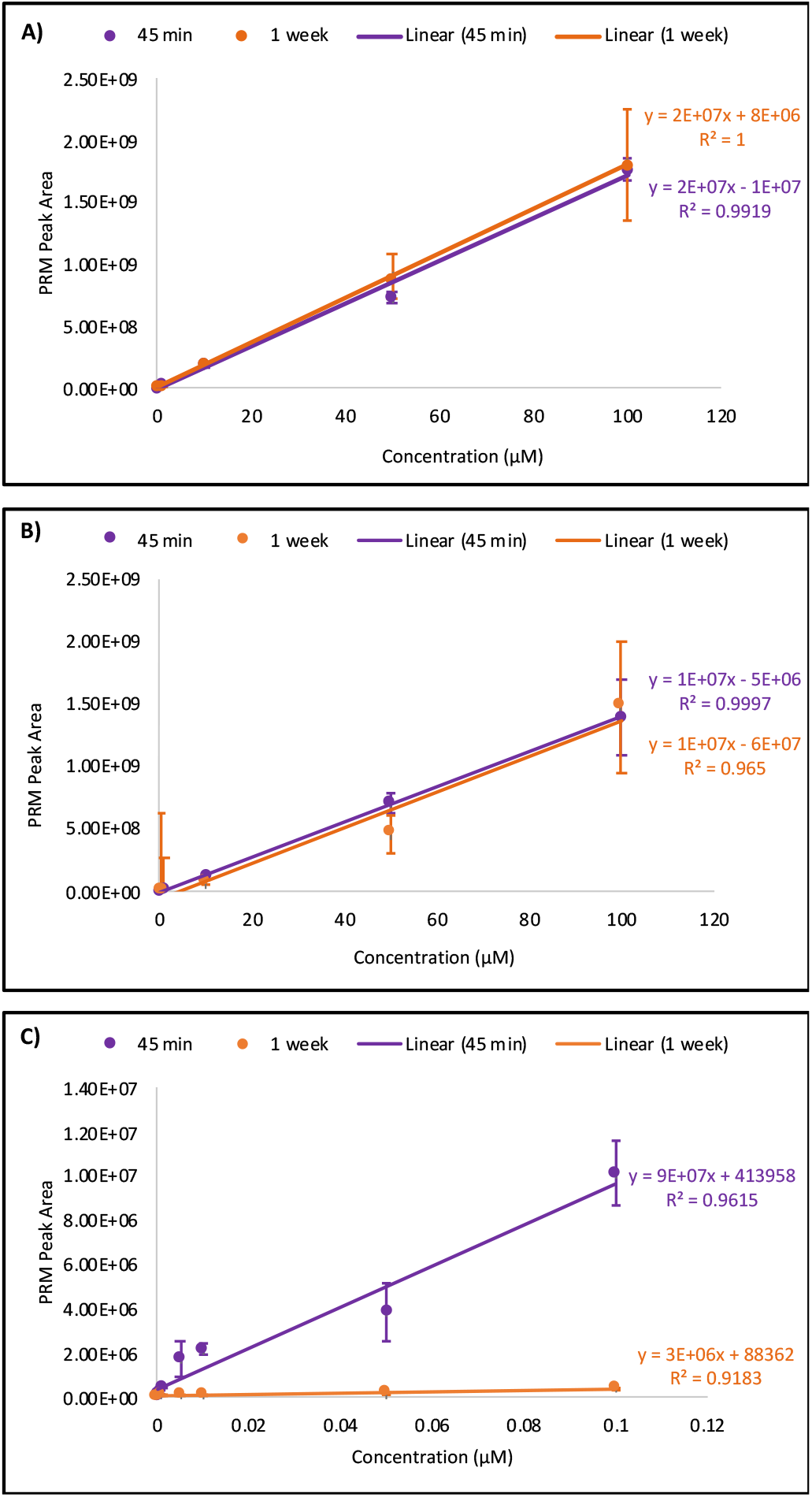
Linearity of swab-based recovery on metal surface at 45 min and 1 week for (A) carnitine, (B) phenylacetylglutamine and (C) cyclobenzaprine. Error bars represent standard error.

### Impact of surface type

In contrast to cyclobenzaprine, which was detected only on one metal surface in our prior built environment analysis ^6a^, carnitine and phenylacetylglutamine were detected on a variety of surfaces, including metal, plastic, glass, cloth, leather and linoleum. Thus, we compared carnitine and phenylacetylglutamine recovery from three additional surface types: linoleum, Plexiglass and a plastified laboratory benchtop surface, in the 0.1-100 μM range, 45 min after compound deposition.

PRM peak area vs spotted concentration was linear for carnitine on plexiglass (R^2^=0.996), benchtop (R^2^=0.9995) and floor (R^2^=0.9581) (**Figure 3**, purple dataset). Likewise, PRM peak area vs spotted concentration was linear for phenylacetylglutamine on plexiglass (R^2^=0.9932), tabletop (R^2^=0.9808) and floor (R^2^=0.9976) (**Figure 4**, purple dataset). Recovery differed between carnitine and phenylacetylglutamine, with minor differences between surface types (16.3%- 27.3% for carnitine vs 10.3%-12.3% for phenylacetylglutamine, **Table 1**), indicating the importance of developing standards for each compound and surface type.

**Figure 3.**
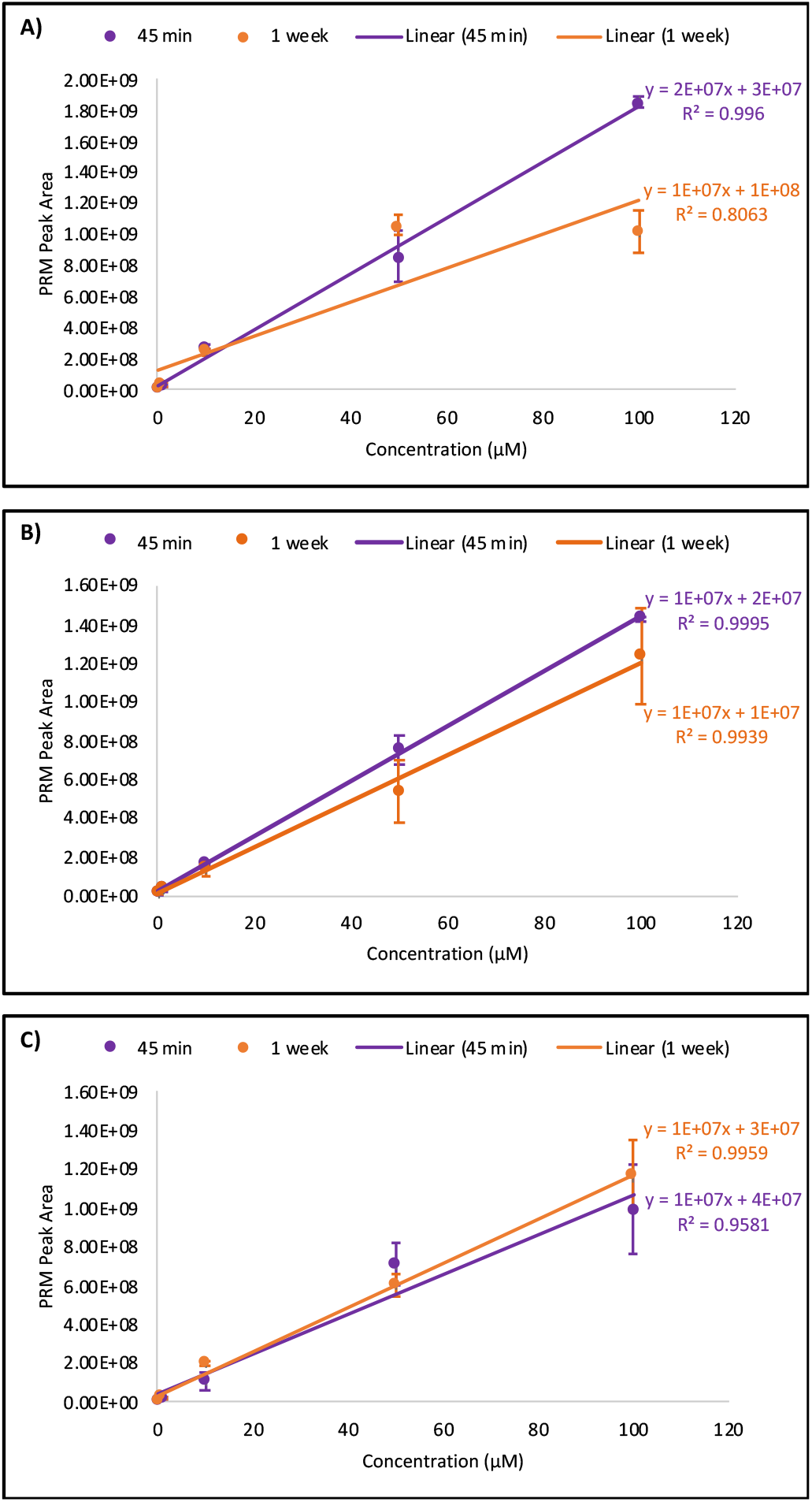
Linearity of swab-based recovery of carnitine on (A) Plexiglass, (B) benchtop and (C) floor surface, 45 min and 1 week after deposition. Error bars represent standard error.

**Figure 4.**
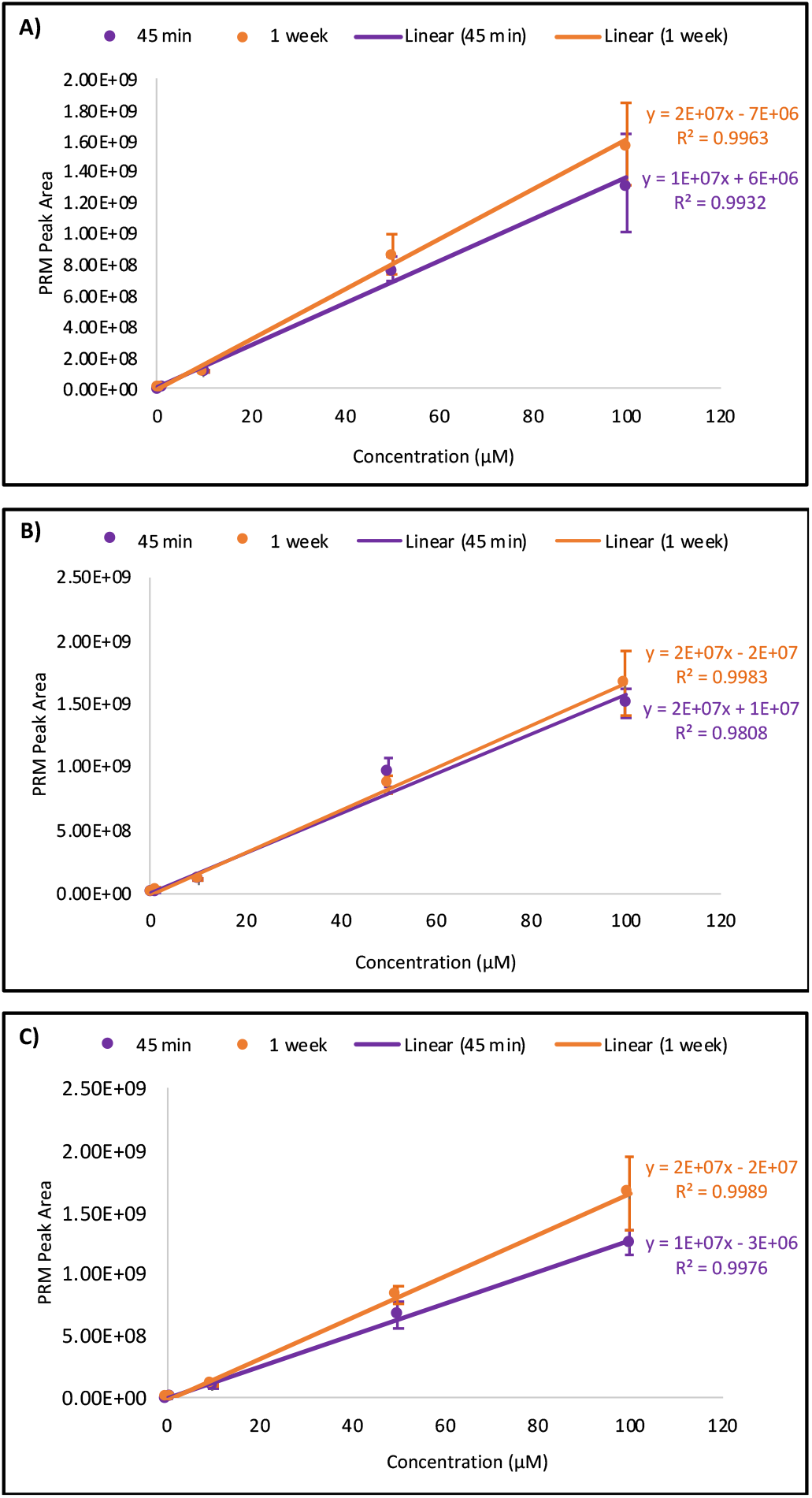
Linearity of swab-based recovery of phenylacetylglutamine on (A) Plexiglass, (B) benchtop and (C) floor surface, 45 min and 1 week after deposition. Error bars represent standard error.

**Table 1.** Average percent recovery (PRM peak area for swab-collected samples compared to pure standards), 45 min post-deposition ± standard error across concentrations.

|  | Metal | Tabletop | Plexi-glass | Floor |
| --- | --- | --- | --- | --- |
| carnitine | 22.6% ± 3.9% | 23.7% ± 2.5% | 27.3% ± 4.1% | 16.3% ± 2.3% |
| phenylacetylglutamine | 10.3% ± 0.7% | 11.9% ± 0.9% | 12.3% ± 1.1% | 10.6% ± 0.6% |

In contrast, oleanolic acid, which had been detected on a variety of surfaces in our prior analyses ^6a^, did not show linearity for swab-based recovery from the plastified benchtop surface, in the 10-100 μM range (**Fig. S1A**, R^2^=0.4413). Oleanolic acid pure standards showed acceptable linearity (**Fig. S1B**, R^2^=0.9094), suggesting instead uneven uptake of surface oleanolic acid by the swab or poor release of oleanolic acid from swab material. A fifth compound, the antibiotic erythromycin, showed poor linearity for pure standards in solution (**Fig. S1C**, R^2^=0.0592) and was not pursued any further.

### Timecourse analysis

In built environment studies, the exact time of chemical deposition is often unknown, and likely to be significantly prior to time of sampling. We therefore assessed whether we could also observe linearity of recovery 1 week post-deposition. For all sampled surfaces and chemicals, we observed similar linearity and recovery slope at 1 week as at 45 min after deposition, except for cyclobenzaprine on metal where recovery was linear but slopes differed between timepoints (**Figure 2, Figure 3, Figure 4**, orange datasets; **Table 2**). Recovery was comparable to what was observed at 45 min post-deposition, with likewise better recovery for carnitine than phenylacetylglutamine (**Table 3**).

**Table 2.** Linear trendline R^2^, 1 week post-deposition.

|  | Metal | Tabletop | Plexi-glass | Floor |
| --- | --- | --- | --- | --- |
| cyclobenzaprine | 0.918<br>3 | ND<sup>a</sup> | ND<sup>a</sup> | ND<sup>a</sup> |
| carnitine | 1 | 0.9939 | 0.8063 | 0.995<br>9 |
| phenylacetylglutamine | 0.965 | 0.9983 | 0.9963 | 0.998<br>9 |

<sup>a</sup> ND, not done: linearity only analyzed on metal surfaces.

**Table 3.** Average percent recovery (PRM peak area for swab-collected samples compared to pure standards), 1 week post-deposition ± standard error across concentrations.

|  | Metal | Tabletop | Plexi-glass | Floor |
| --- | --- | --- | --- | --- |
| carnitine | 23.6% ± 4.4% | 24.7% ± 3.5% | 22.6% ± 2.6% | 18.5% ± 2.8% |
| phenylacetylglutamine | 8.2% ± 0.8% | 9.7% ± 1.1% | 12.6% ± 1.1% | 11.8% ± 0.7% |

### Implementation

To address the utility of this method in estimating chemical abundance and transfer from built environment surfaces, we used the same method to sample from metal surfaces in the workplace built environment. Observed peak area for carnitine was 10,595,825 on a door handle and 10,024,754 on a water fountain button. Using the standard curves for long-term carnitine recovery from metal (**Figure 3A**), we estimate a recovered concentration of 0.136 μM and 0.104 μM for each of these surfaces, respectively.

## CONCLUSIONS

Overall, we have developed a straightforward and affordable method to quantitatively assess surface chemical levels. This method can be applied to a broad range of chemicals of interest to the exposome and behavioral fields, including the following Classyfire classes: organonitrogen compounds (Classyfire subclass: quaternary ammonium salts, carnitine), carboxylic acids (Classyfire direct parent: N-acyl-alpha amino acids, phenylacetylglutamine) and dibenzocycloheptenes (cyclobenzaprine) ^9^. One limitation however is that it is not suited to all chemicals or surface types. Thus, linearity of recovery should be assessed empirically for chemicals of interest, and standard curves built for each specific chemical and surface of interest, with time-dependence of signals assessed. By applying proper precautions when building reference standards on surfaces, for example by building standard curves in a fume hood, this method could readily be applied to hazardous chemicals. Likewise, for non-toxic chemicals, dilution references could be built on human skin.

We acknowledge that our method is unable to fully recover surface chemicals (recovery of 8.2%-27.3% compared to the deposited chemical concentration). Although this can be addressed by determining the recovery factor to extrapolate the actual surface chemical concentration, more importantly, our method actually assesses the amount of chemicals that can be transferred from this surface. It therefore represents a more relevant measurement in the context of chemical exposure assessment. We acknowledge that the recovered measured peak area may be below the linear quantification range for some chemicals. However, we were able to estimate concentrations for chemicals of interest in this study. We therefore anticipate this method to have broad applicability in exposomics and human behavior studies.

## ASSOCIATED CONTENT

### Supporting Information

Supplementary table ST-1. Skyline parameters.

Supplementary Figure S1. Metabolites unsuitable for swab-based quantitative analysis. (A) Swab-based recovery of oleanolic acid from laboratory benchtop. (B) Oleanolic acid pure standard in solution. (C) Erythromycin pure standard in solution.

## AUTHOR INFORMATION

### Author Contributions

The manuscript was written through contributions of all authors. / All authors have given approval to the final version of the manuscript. /

## ACKNOWLEDGMENT

This project was supported by start-up funds from the University of Oklahoma.

## Notes

### Competing Interest Statement

The authors have declared no competing interest.

