## Supplementary figures and images for "Optimized extraction method enables quantitative analysis of surface metabolite recovery for exposomics and behavioral studies"

### Figure S1

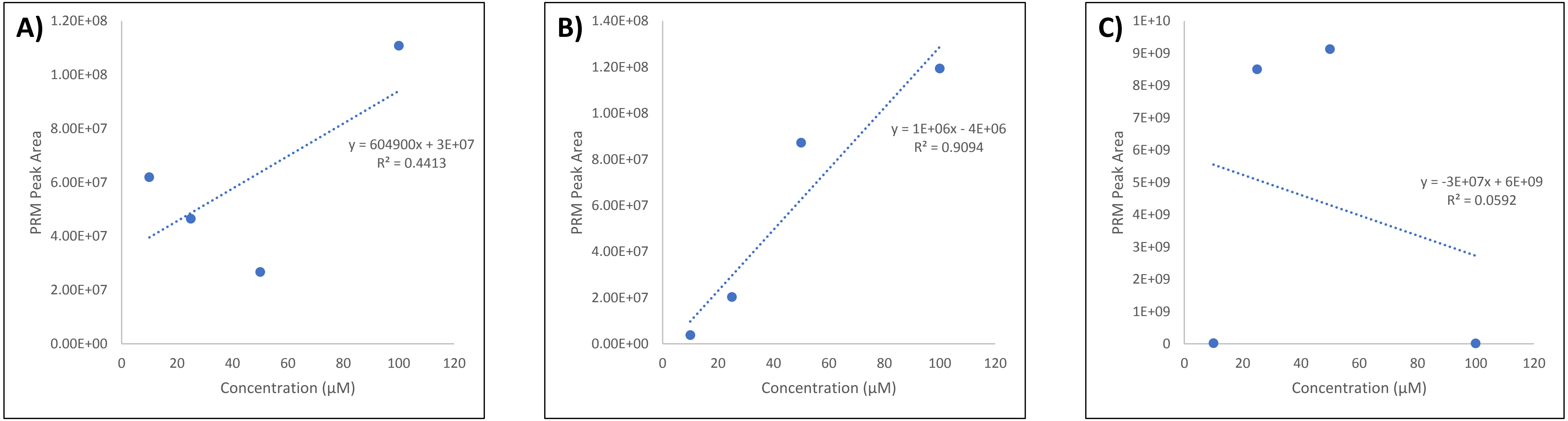
